# Gene-experience correlation during cognitive development: Evidence from the Adolescent Brain Cognitive Development (ABCD) Study^SM^

**DOI:** 10.1101/637512

**Authors:** Robert J. Loughnan, Clare E. Palmer, Wesley K. Thompson, Anders M. Dale, Terry L. Jernigan, Chun Chieh Fan

**Author notes:** Shared Corresponding, equal contribution.

## Abstract

**Background:** Findings in adults have shown more culturally sensitive ‘crystallized’ measures of intelligence have greater heritability, these results were not able to be shown in children.

**Methods:** With data from 8,518 participants, aged 9 to 11, from the Adolescent Brain Cognitive Development (ABCD) Study^®^, we used polygenic predictors of intelligence test performance (based on genome-wide association meta-analyses of data from 269,867 individuals) and of educational attainment (based on data from 1.1 million individuals), associating these predictors with neurocognitive performance. We then assessed the extent of mediation of these associations by a measure of recreational reading.

**Results:** more culturally sensitive ‘crystallized’ measures were more strongly associated with the polygenic predictors than were less culturally sensitive ‘fluid’ measures. This mirrored heritability differences reported previously in adults and suggests similar associations in children. Recreational reading more strongly statistically mediated the genetic associations with crystallized than those with fluid measures of cognition.

**Conclusion:** This is consistent with a prominent role of gene-environment correlation in cognitive development measured by “crystallized” intelligence tests. Such experiential mediators may represent malleable targets for improving cognitive outcomes.

## Introduction

Scores on cognitive tests in both children and adults have been linked to long term outcomes and to genetic variation(1–4). Some cognitive tests, e.g., those requiring literacy and mathematical skills, depend upon and are more sensitive to variability in cultural and socio-economic factors. These measures are often referred to as ‘crystallized’ intelligence measures. In contrast, other tests that tap the capacity to solve novel problems, or process novel information, often referred to as ‘fluid’ measures, are less culturally sensitive and are less strongly related to socio-economic variables(5,6). A recent review reported systematic differences in heritability (an estimate of trait variability attributable to genetic variation) of the traits measured by these different kinds of cognitive measures(7). Surprisingly, in studies of adult twins, more culturally sensitive tests exhibited higher, rather than lower, heritability; which runs counter to predictions from conventional models of intelligence. The authors described similar trends in the twin studies of children, but increased heritability of crystallized relative to fluid measures have not yet been established for children, in whom intellectual functions are continuing to mature.

The finding that the measures most strongly influenced by cultural factors exhibit higher heritability is perhaps counterintuitive; however previous authors have noted that genetic variation can be associated with environmental, cultural, or experiential (ECE) factors that themselves amplify effects of a genotype on the phenotype, a phenomenon often referred to as rGE (gene-environment correlation). These associations between genotypes and ECE factors could influence the development of cognitive and intellectual abilities in several ways. As an example, if others in the social environments of children recognize traits, e.g., precocious behavior, in those with a genetic propensity for a given cognitive ability, they may begin to treat such individuals differently, rewarding them disproportionately for intellectual pursuits, investing more in their instruction, and/or placing them in environments that drive learning more effectively. Alternatively, the associations can be driven by the motivation of the children themselves if for example they develop greater enthusiasm for intellectual activities for which they have been more frequently rewarded, and which they then pursue more assiduously, thus enjoying beneficial effects of the increased practice associated with these activities. In either case, the genetically advantaged abilities are disproportionately enhanced by these mediating ECE factors. Of course, individuals with less advantageous genotypes may experience the converse of these social and motivational effects, resulting in languishing, or in the worst case suppressed, intellectual development, even within similar environments. Such rGE effects can increase variance in intellectual phenotypes and increase estimates of heritability using both epidemiological and genomic methods(8). The important implication is that a component of this increased heritability requires the mediating ECE effects for its expression. In essence, more direct biological effects of the genotype *and* associated differences in the environments or experiences of the child are both contributing causal factors influencing the mature phenotype, but they act through dissociable mechanisms.

Heritability is a population statistic frequently measured using a twin design. For this study, we used polygenic scores to examine variation in genetic and experiential factors and their relationship to trait measures of cognitive function. Polygenic scores have the advantage that they can be used to index relevant genetic factors in samples of unrelated individuals by leveraging the statistical power of meta-analysis results from large Genome Wide Association Studies (GWAS). Using neurocognitive test scores, genomic data, and a measure of parent-reported recreational reading assessed in a large sample of 8,618 children, aged 9 to11, from the ABCD Study®, we used polygenic scores of intelligence test performance (based on GWAS of 269,867 individuals(9)) and educational attainment, sometimes considered a proxy for intellectual ability (based on 1.1 million individuals(10)), to ask 3 questions: First, do these genomic predictors account for more of the variability in estimates of culturally sensitive crystallized traits than fluid traits in children, as might be expected from reports of higher heritability in adult twins? Second, does a parent-reported estimate of the time their children spend reading for pleasure mediate the relationship between a genomic predictor and measures of cognitive performance, consistent with a role of this experiential enhancer of performance in increasing heritability? Third, if mediation is observed, is this mediating effect larger for the culturally sensitive crystallized than the fluid measures of cognitive performance, consistent with a role for rGE in the higher heritability of these measures?

In additional analyses, we examined the degree to which the findings in the ethnically diverse ABCD sample were similar between the subgroup of children with high genomic European ancestry (EurA) and a remaining subgroup of children who were from diverse ancestry groups (DivA). Finally, using simulations, we tested whether our observed findings may be due to previously reported differences in test-retest reliabilities (for crystallized vs fluid measures).

## Materials and Methods

### 2.1 Data available in the ABCD data release 2.0.1

The ABCD study (http://abcdstudy.org) enrolled the families of 11,875 children aged 9 or 10 years at baseline(11). This longitudinal study follows the development of these children at 21 sites across the US for ten years. The cohort exhibits a large degree of sociodemographic diversity. Exclusion criteria were limited to: 1) lack of English proficiency; 2) the presence of severe sensory, neurological, medical or intellectual limitations that would inhibit the child’s ability to comply with the protocol; 3) an inability to complete an MRI scan at baseline. The study protocols are approved by the University of California, San Diego Institutional Review Board(12). Parent/caregiver permission and child assent from each participant were obtained. Here, our data were drawn from the baseline assessments shared in ABCD release 2.0.1 (NDAR DOI: 10.15154/1504041).

#### 2.1.1 Cognitive Measures

Seven of the 10 cognitive tasks were subtests from The NIH Toolbox Cognition Battery^®^ (NTCB) in the version recommended for ages 7+(http://www.nihtoolbox.org)(13). The average time to complete this battery is approximately 35 minutes. The NTCB was administered in English(14), using an iPad, with support from a research assistant when needed. The battery yields individual test scores measuring specific constructs and composite scores that have been shown to be highly correlated with ‘gold standard’ measures of intelligence in adults(15) and children(5). Here, all 7 individual test scores and 2 composite scores were examined: the Crystallized Cognition Composite Score (derived from scores on the Picture Vocabulary and Oral Reading Recognition measures) and the Fluid Cognition Composite Score (derived from the remaining measures). Additionally, three neurocognitive tasks were used that were not components of the NTCB: Rey-Auditory Learning Task, Little Man Task and Matrix Reasoning. Please see supplementary materials for a description of each task.

#### 2.1.2 Latent Neurocognitive Factors

Thompson et al. derived a three factor, varimax rotated, solution for the latent structure across the neurocognitive battery in ABCD using Bayesian Probabilistic PCA(16). The final latent factor solution included the measures described above, except for the Matrix Reasoning task which had very little effect on the solution. The factors will be referred to as Bayesian Factors (BF) 1-3. Language tasks loaded most heavily on BF1, which was highly correlated with the Crystallized Composite (r=0.93); executive functioning tasks loaded most heavily on BF2; and learning/memory tasks loaded heavily on BF3.

#### 2.1.3 Recreational Reading

Parents of ABCD participants were asked to complete a survey of their children’s activities. One question asked, “Does your child read for pleasure?” The follow-up question was, “About how many hours per week does your child read for pleasure?”. This estimate of number of hours of recreational reading was log transformed due to skewness. To confine the analyses of this variable to a homogenous group of children who read for pleasure, we included only children whose parents answered ‘yes’ to the first question.

#### 2.1.4 Genetic Data and Computing Polygenic Scores

Using genotype data we derived genetic ancestry using fastStructure(17) with four ancestry groups. Genetic principal components were also calculated using PLINK. Variants were imputed using the Michigan Imputation Server(18). Polygenic scores were computed using PRSice(19). The Intelligence Polygenic Score (IPS) was trained on 269,867 individuals by Savage et al.(9), and focused on neurocognitive tests considered to gauge fluid intelligence. The Education Attainment Polygenic Score (EAPS) was generated from 1.1 million individuals, predicting the phenotype of number of years of schooling completed. Please see supplementary materials for further details on genetic data and analysis.

We were primarily focused on studying the IPS association with cognitive tests in ABCD, due to it being trained on a more directly relevant phenotype. However, we additionally examined EAPS as a secondary analysis for comparison as it has been previously used as a proxy for cognitive ability and has a discovery sample size four times the size of the IPS.

### 2.2 Analytic Methods

#### 2.2.1 Ancestry Group Analyses

Training and testing polygenic scores in different ancestry groups has been shown to reduce predictive power(20–22). Given the ancestry differences between the polygenic score discovery samples (predominantly European) and the ABCD study (multiple ancestry groups), we wanted to confirm our main results in the full samples were not driven by population structure. As such we additionally performed analyses in two subsamples: 1) children with estimated proportion of European ancestry higher than 90% (EurA) and 2) a group of the remaining children with diverse ancestry, which included those from other or mixed ancestry (DivA).

#### 2.2.2 Statistical Model for Genomic Prediction of Behavioral Measures

To assess the association between the polygenic scores and cognitive performance in ABCD, we fit Generalized Linear Mixed-Effects Models (GLMMs) using the gamm4 package(23) in R. Each model predicted performance on a different cognitive measure. Continuous variables were z-scored before model fitting to allow coefficients to be interpreted as standardized effect sizes. To test if regression coefficients differed between regressions we performed a z-test on the difference between coefficients, based on the propagated standard error for the two regression coefficients as the sum of the error of variances for each measure. This test assumes the standard errors are uncorrelated and so provides a conservative estimate of significance. Please see supplementary materials for details and covariates used.

#### 2.2.3 Differential Mediation Analysis

To assess whether recreational reading is a plausible ECE factor increasing heritability of crystallized cognition, through rGE effects, we performed a mediation analysis. Specifically, we compared the statistical mediation effects of recreational reading experience on the associations between the IPS and both the Crystallized Composite and Fluid Composite, respectively. We achieved this by calculating the proportion of mediation of recreational reading on the i) IPS-crystallized and the ii) IPS-fluid associations using an average causal mediation effect model(24) across 10,000 bootstrap samples. With bootstrapped samples we tested if the mediation effect of recreational reading on the IPS-crystallized association was greater than that of IPS-fluid, by performing a Welch t-test on the samples. Mediation analysis was performed using general linear models in the mediation package in R(25), see supplementary materials for details.

#### 2.2.4 Data and Code Availability

The ABCD dataset is available approved researchers at https://nda.nih.gov/abcd. A jupyter notebook of the analysis be found at github.com/robloughnan/ABCD_Intelligence_Polygenic_Score.

## Results

### Demographics

Figure *1* illustrates a flow-chart for sample selection. For the final analysis we have 8,518 individuals in the full sample, 4,885 in the EurA sample and 3,633 in the DivA sample.

**Figure 1.**
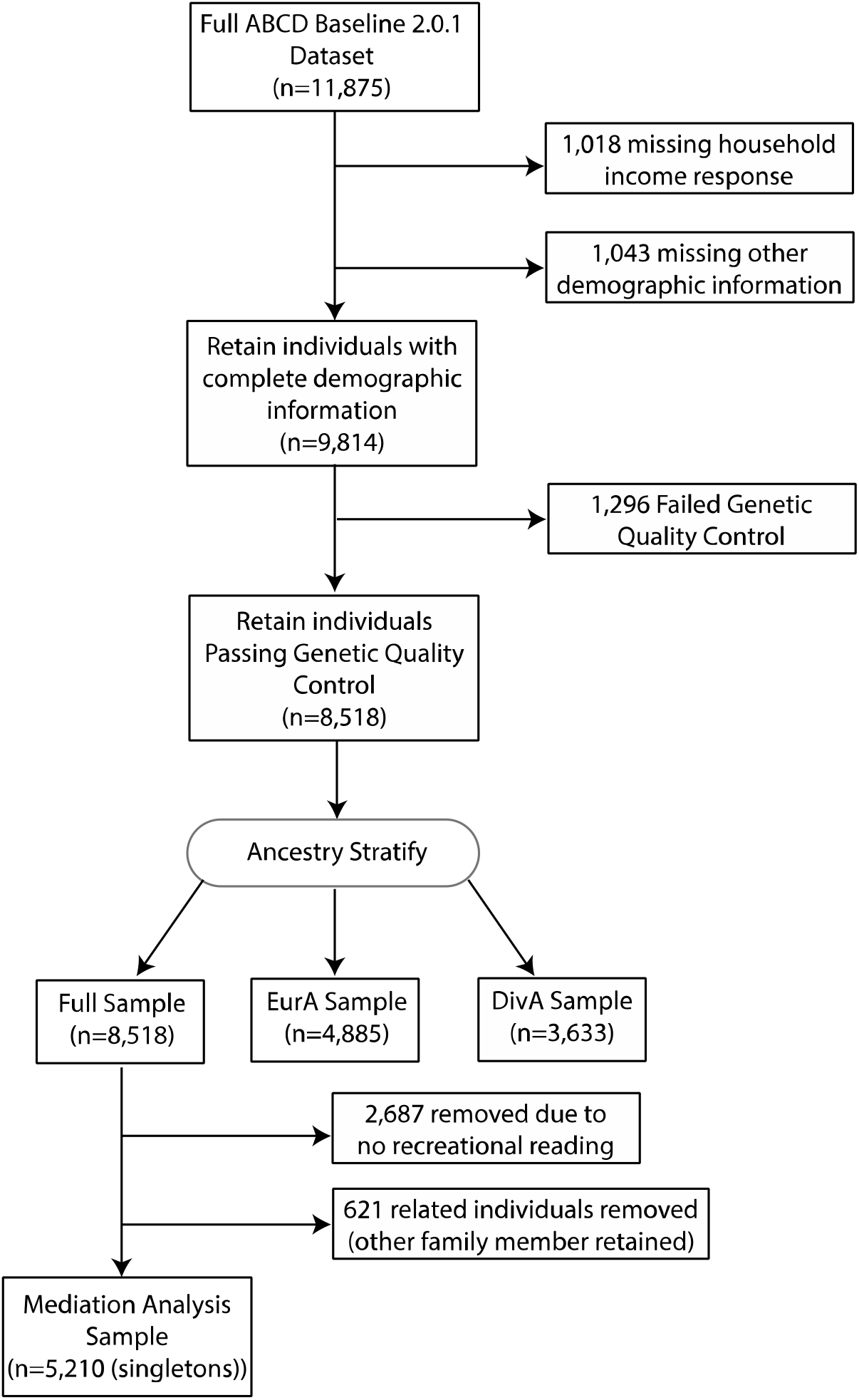
Flow chart of sample selection and exclusion.

### Behavioral Measures and Sociocultural Factors

Mean performance, standard deviation (SD), median and estimates of variance explained by age, sex, and the set of socio-cultural covariates (parental marital status, highest education level of parent/caregiver, household income, ethnicity, genetic principal components) are given for each behavioral measure examined in Table 2. Consistent with previous reports, there are substantial differences in the degree to which socio-cultural factors account for variability in these measures. The Crystallized Composite, its constituent Picture Vocabulary and Reading Recognition measures, and BF1, on which these measures of language and literacy load heavily, all exhibit higher levels of association with socio-cultural variables. This pattern persisted when controlling for IPS (Sup. Table 2). Sex, age and socio-cultural factors explained little variability in recreational reading. Partial correlations between the individual cognitive task measures controlling for covariates (Figure 2), suggest that performance on the different tasks is modestly correlated across children (rs=.08-.41) in this sample. Correlations peak in the.3 range within Fluid Composite measures, and the highest correlation is observed between the two Crystallized Composite measures (Picture Vocabulary and Oral Reading r=.41).

**Table 1:** Summary of demographics for individuals included in the full sample for the present genomic prediction analyses, and for the genomic European Ancestry and genomic Other Ancestry subgroups.

|  | *Full Sample* | *European Ancestry* | *Other Ancestry* |
| --- | --- | --- | --- |
| **Total N** | 8518 | 4885 | 3633 |
|  | **Mean (SD)** |  |  |
| *Age - months* | 119.05 (7.48) | 119.21 (7.49) | 118.85 (7.47) |
|  | **N (%)** |  |  |
| *Sex Male* | 4438 (52.1) | 2576 (52.7) | 1862 (51.3) |
| *Parent Married = Yes* | 6024 (70.7) | 4066 (83.2) | 1958 (53.9) |
| **Parental Education** |  |  |  |
| *< HS Diploma* | 302 (3.5) | 21 (0.4) | 281 (7.7) |
| *HS Diploma/GED* | 649 (7.6) | 138 (2.8) | 511 (14.1) |
| *Some College* | 2149 (25.2) | 899 (18.4) | 1250 (34.4) |
| *Bachelor* | 2318 (27.2) | 1548 (31.7) | 770 (21.2) |
| *Post Graduate Degree* | 3100 (36.4) | 2279 (46.7) | 821 (22.6) |
| **Household Income** |  |  |  |
| *[<50K]* | 2353 (27.6) | 596 (12.2) | 1757 (48.4) |
| *[>=50K & <100K]* | 2444 (28.7) | 1471 (30.1) | 973 (26.8) |
| *[>=100K]* | 3721 (43.7) | 2818 (57.7) | 903 (24.9) |
| **Race** |  |  |  |
| *White* | 5715 (67.7) | 4750 (97.4) | 965 (27.1) |
| *Black* | 1129 (13.4) | 1 (0.0) | 1128 (31.7) |
| *Asian* | 199 (2.4) | 0 (0.0) | 199 (5.6) |
| *Other* | 1397 (16.6) | 128 (2.6) | 1269 (35.6) |
| **Hispanic** |  |  |  |
| *Hispanic* | 1628 (19.1) | 131 (2.7) | 1497 (41.2) |

**Table 2.**
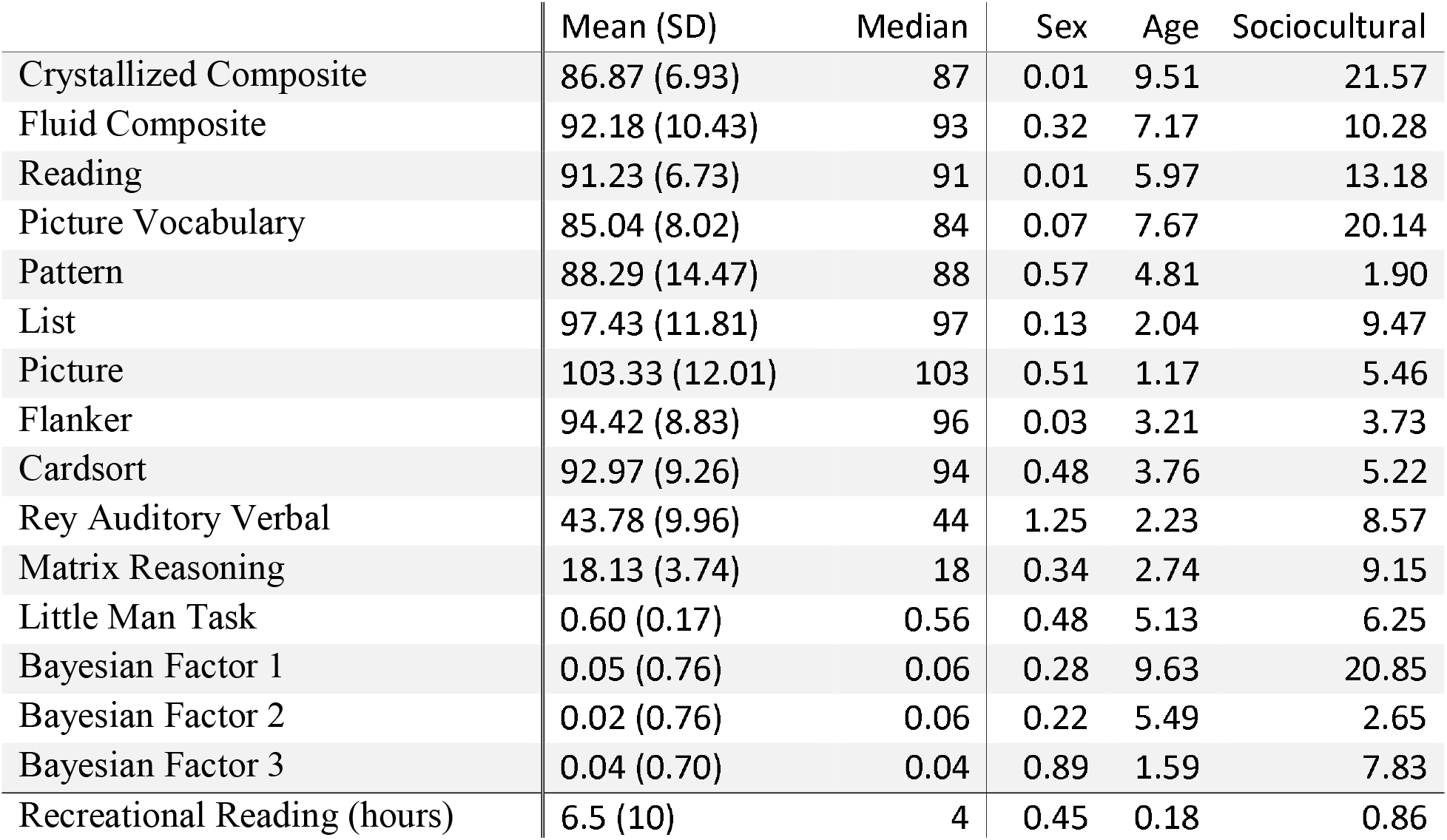
Mean (SD) and median for each behavioral measure in the full sample, estimated % variance explained by sex, age, and the set of socio-cultural covariates (parental marital status, parental education, household income, genetic ancestry PCs and Hispanic/non-Hispanic).

|  | Mean (SD) | Median | Sex | Age | Sociocultural |
| --- | --- | --- | --- | --- | --- |
| Crystallized Composite | 86.87 (6.93) | 87 | 0.01 | 9.51 | 21.57 |
| Fluid Composite | 92.18 (10.43) | 93 | 0.32 | 7.17 | 10.28 |
| Reading | 91.23 (6.73) | 91 | 0.01 | 5.97 | 13.18 |
| Picture Vocabulary | 85.04 (8.02) | 84 | 0.07 | 7.67 | 20.14 |
| Pattern | 88.29 (14.47) | 88 | 0.57 | 4.81 | 1.90 |
| List | 97.43 (11.81) | 97 | 0.13 | 2.04 | 9.47 |
| Picture | 103.33 (12.01) | 103 | 0.51 | 1.17 | 5.46 |
| Flanker | 94.42 (8.83) | 96 | 0.03 | 3.21 | 3.73 |
| Cardsort | 92.97 (9.26) | 94 | 0.48 | 3.76 | 5.22 |
| Rey Auditory Verbal | 43.78 (9.96) | 44 | 1.25 | 2.23 | 8.57 |
| Matrix Reasoning | 18.13 (3.74) | 18 | 0.34 | 2.74 | 9.15 |
| Little Man Task | 0.60 (0.17) | 0.56 | 0.48 | 5.13 | 6.25 |
| Bayesian Factor 1 | 0.05 (0.76) | 0.06 | 0.28 | 9.63 | 20.85 |
| Bayesian Factor 2 | 0.02 (0.76) | 0.06 | 0.22 | 5.49 | 2.65 |
| Bayesian Factor 3 | 0.04 (0.70) | 0.04 | 0.89 | 1.59 | 7.83 |
| Recreational Reading (hours) | 6.5 (10) | 4 | 0.45 | 0.18 | 0.86 |

**Figure 2.**
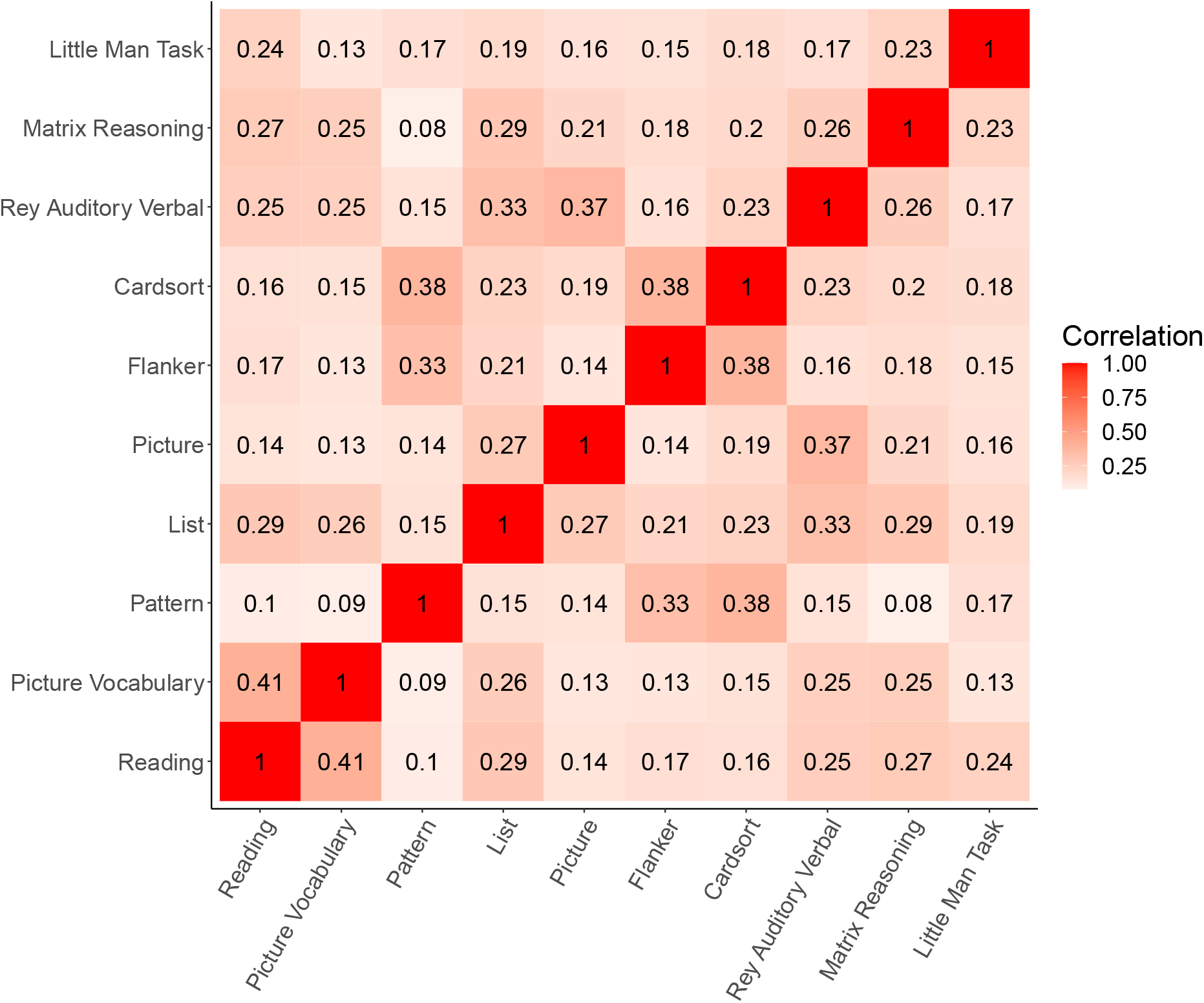
Partial correlation matrix showing intercorrelations among individual task performance measures (controlling for age, sex, parental marital status, parental education, household income, principal components of genetic ancestry and Hispanic status) in the full sample included in the present study of genomic predictors.

### Genomic Prediction of Crystallized and Fluid Cognition Measures

Table 3 summarizes the regression results for predicting the Crystallized and Fluid Composites with IPS or EAPS in the full sample, and separately in the EurA and DivA subsamples. The IPS was a significant predictor of both measures in all analyses. Importantly, the standardized regression coefficient was significantly higher for the Crystallized than the Fluid Composite regardless of ancestry group (full sample: z=4.8, p=1.8×10^−6^, EurA: z=4.6, p=5.1×10^−6^ and DivA: z=2.5, p=1.4×10^−2^).

**Table 3.** Regression results for GLMMs associating IPS (top) and EAPS (bottom) with Crystallized Composite and Fluid Composite of the NIH toolbox within full Sample and ancestry subgroups.

| Sample | Fluid Composite |  |  |  | Crystallized Composite |  |  |  |
| --- | --- | --- | --- | --- | --- | --- | --- | --- |
| | Stand. $\beta$ | t | P value | % Var. Explained | Stand. $\beta$ | t | P value | % Var Explained |
|  | IPS |  |  |  |  |  |  |  |
| Full Sample | 0.28 | 8.03 | 1.14E-15 | 0.75 | 0.50 | 15.82 | 1.31E-55 | 2.86 |
| EurA | 0.11 | 7.53 | 6.10E-14 | 1.15 | 0.21 | 14.48 | 1.44E-46 | 4.13 |
| DivA | 0.20 | 3.41 | 6.52E-04 | 0.32 | 0.40 | 7.34 | 2.68E-13 | 1.47 |
|  | EAPS |  |  |  |  |  |  |  |
| Full Sample | 0.11 | 7.23 | 5.26E-13 | 0.61 | 0.19 | 14.21 | 2.56E-45 | 2.32 |
| EurA | 0.09 | 6.60 | 4.66E-11 | 0.89 | 0.18 | 12.95 | 9.36E-38 | 3.34 |
| DivA | 0.08 | 3.38 | 7.28E-04 | 0.32 | 0.15 | 6.66 | 3.24E-11 | 1.21 |

In no case did the EAPS, despite a much larger training sample size, appear to account for more of the variance in the neurocognitive measures than did IPS. However, across ancestry groups and for both composite scores, combining both genomic predictors explained significantly more variance in behavior than IPS alone (supplementary results). IPS + EAPS explained 5.8% variance (p=4.5×10^−64^) in the Crystallized Composite for EurA (a 40% increase compared to IPS alone). Supplementary Tables 3-8 show regression results for each behavior using IPS, EAPS and IPS + EAPS within each ancestry group.

Fitting separate regression models for each individual task in the neurocognitive battery, we found that the IPS was a significant predictor for each cognitive measure for the full sample and the EurA subsample (all p values<10^−3^), surviving the Bonferroni-corrected significance threshold of 0.05/10=0.005. Within the DivA subsample only six of the ten tasks were individually significantly predicted by the IPS (Sup. Table 8). Figure 3 shows the standardized regression coefficients of IPS predicting performance on each task, as well as the Crystallized and Fluid Composite measures from the NTCB and Bayesian Factors 1-3(16), in the full sample. Individual cognitive measures included in the Crystallized Composite have consistently higher IPS standardized regression weights than the measures included in the Fluid Composite. Other neurocognitive tasks from the ABCD battery (shaded in gray) showed similar associations to the Fluid Composite. The results for the Bayesian Factors mirrored these results: BF1, on which ‘Crystallized” measures had the highest factor loadings (Sup. Figure 1)(16), displayed a stronger association with IPS than BF2 and BF3 on which ‘Fluid”, executive function and memory measures had higher loadings. The results in the subsamples (EurA and DivA) are provided in supplemental material.

**Figure 3.**
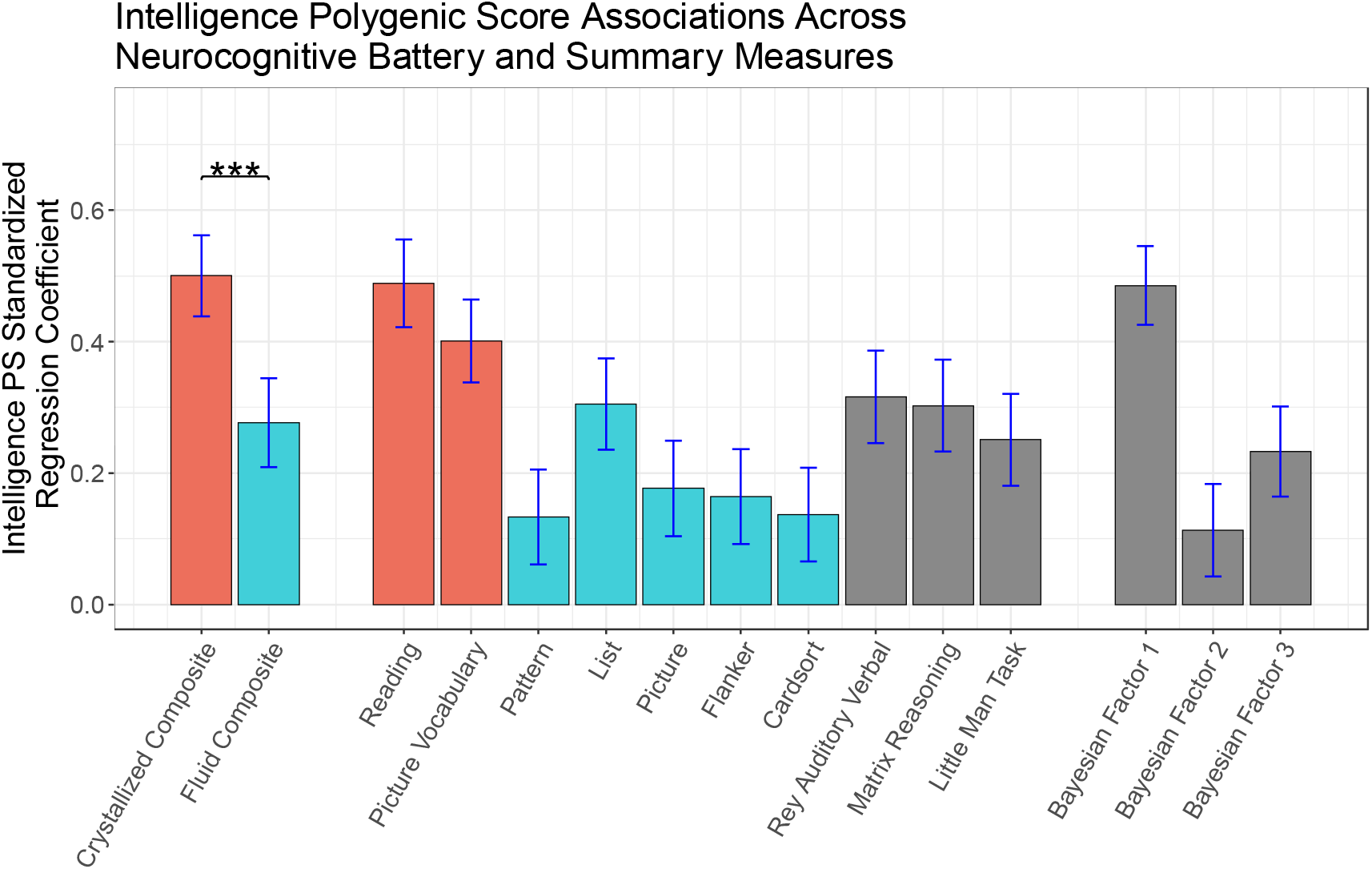
Standardized regression coefficients of IPS for fitting linear mixed models to performance on Fluid and Crystallized Composites, each individual task from the NTCB, additional measures from the ABCD neurocognitive battery, and Bayesian (latent) Factors 1-3, in the full sample. Prediction of the Crystallized Composite is significantly stronger than for the Fluid Composite. Tasks included in the Fluid Composite (shaded in blue) have consistently lower regression coefficients than those included in the Crystallized Composite (shaded in red). Additional measures from the neurocognitive battery exhibit associations with IPS more similar to the Fluid Composite than to the Crystallized Composite, however Bayesian Factor 1, on which the verbal tasks load heavily, exhibits an association similar to the Crystallized Composite. Error bars show estimates of 95% confidence intervals as 1.96 × standard error.

### Differential Mediation Results

The mediation analysis showed that recreational reading partially mediated associations between IPS and both composite measures, proportions of mediation: fluid 0.084 (95% CI: 0.047-0.14, p≤10^−4^), crystallized 0.12 (95% CI: 0.88-0.16, p≤10^−4^). However, the differential mediation analyses revealed a highly significant difference between the large degree of attenuation of the association between IPS and the Crystallized Composite relative to that between IPS and the Fluid Composite (Welch t-test: t=125, df=19053, p<10^−300^), shown in Figure 4.

**Figure 4.**
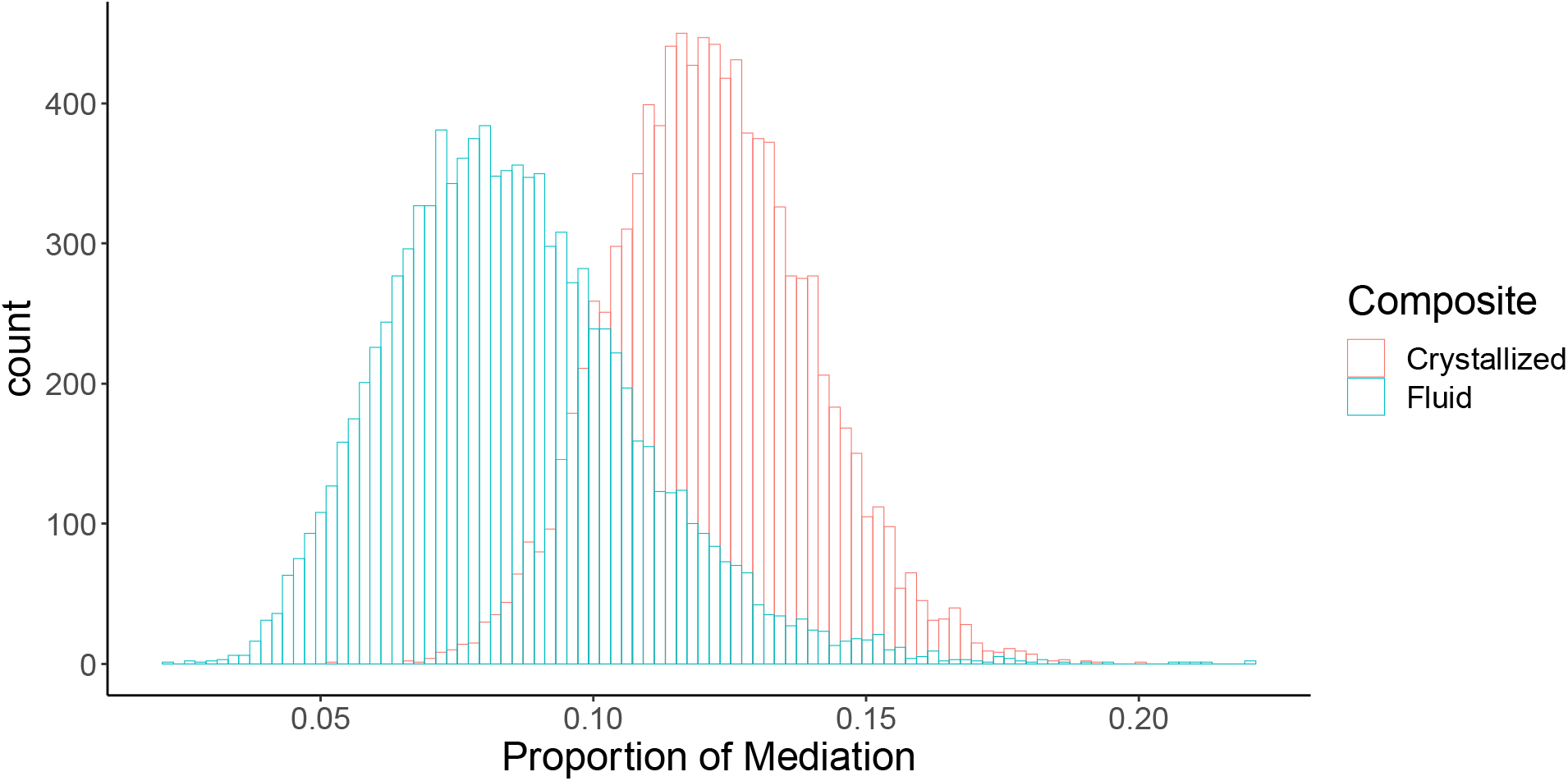
Differential mediation analysis in singletons (N= 5,210): histograms shows 10,000 bootstrap estimates for proportion of mediation of recreational reading on: i) IPS and Crystallized Composite (red) and ii) IPS and Fluid Composite (blue). Recreational reading attenuates the relationship between the IPS and the Crystallized Composite to a significantly greater degree.

### Sensitivity Analyses to Address Test Reliability

A previous study reported the test-retest reliability for the Fluid Composite from the NTCB (.76) was somewhat lower than for the Crystallized Composite (.85)(5), raising questions about whether differences in the strength of their associations with IPS could be attributed to more noise in the Fluid Composite measure. In supplementary sensitivity analyses we demonstrate that our results are robust to the addition of simulated noise to the Crystallized Composite that mimics this difference in test reliability. At this level of simulated noise we estimated 1.0 power (alpha=0.05) to detect *Cryst_noise_* having a significantly greater IPS standardized regression coefficient than the Fluid Composite. Moreover, additional sensitivity analyses indicate that the observed differences in the mediation effects of recreational reading are similarly robust against potential measurement error modelled as random noise. These analyses are described in detail in Supplemental Material.

## Discussion

We have shown that polygenic predictors of intelligence test performance and of educational attainment are associated with neurocognitive performance in this large group of children from diverse backgrounds. These results are consistent with previous findings demonstrating that virtually all behavioral traits, including cognitive and intellectual phenotypes, are heritable(26). Moderate estimates of heritability of many behavioral phenotypes also establish that a substantial portion of the variability is due to independent environmental influences. Given that behavioral phenotypes emerge through interactions between children and their physical, social, and cultural environments, much attention has been paid to how these environmental factors modify the phenotypes, since they are presumably the malleable factors. However, recently, more attention has been focused on the possible roles of mediating nongenetic (ECE) factors that, through their statistical association with genetic variation (rGE), may amplify heritability(7,8).

We found that a culturally dependent estimate of crystallized cognitive functions, the Crystallized Composite measure from the NTCB, is more strongly associated with the best available polygenic predictor of intelligence test performance than is the Fluid Composite measure, consistent with earlier findings in adults of heritability differences(7) and polygenic score performance(27) across similar measures. This is despite the IPS being based on a large meta-analysis of GWAS combining cognitive measures that were described by the authors as primarily “fluid intelligence” measures(9). Indeed, the relative size of the IPS association across the 15 measures examined here (Figure 2) closely mirrored the relative percent variance explained in these measures by socio-cultural variables (Table 2), a pattern that persists after accounting for IPS (Sup. Table2). Moreover, for children who read for pleasure, the extent of recreational reading was found to partially mediate the associations between IPS and both Composite measures, but to a significantly greater degree for the crystallized than for the fluid measure, consistent with a more prominent role of rGE in the development of abilities tapped by measures that are both more heritable, and apparently more sensitive to socio-cultural variables. In other words, even when controlling for *independent* contributions of more global sociocultural variables, how often a child reads for pleasure more strongly mediates the association between IPS and crystallized rather than in fluid performance.

It is perhaps unsurprising that recreational reading more strongly mediates the association between IPS and culturally sensitive measures of intelligence since such measures are generally sensitive to educational factors. Indeed, a measure of oral reading proficiency loads highly on both measures of crystallized functions examined here, Crystallized Composite and BF1. One can imagine that children with neurobehavioral phenotypes advantageous for learning to read might be more likely to develop the habit of reading for pleasure than those with other neurobehavioral phenotypes, for a variety of reasons. However, these results imply that choosing to read for pleasure at 10 years of age is associated with having a genotype linked to intellectual functions most dependent on reading, and that an estimate of the frequency of reading behavior mediates that link. This is consistent with previous descriptions of rGE effects, and with analyses by Beam and Turkheimer(8), who showed that increasing rGE over time could explain observed increases in the heritability of measures of cognitive function through development. The ABCD study will provide an opportunity to measure changes in heritability at later time points of this longitudinal study. Importantly, despite the lower test-retest reliability of the fluid compared to the crystallized composite score from the NTCB(5), our supplementary analyses show that this difference in test reliability is unlikely to explain our findings.

Though recreational reading would appear to be an enhancing mediator of intellectual development, it is important to note that genotype-correlated ECE factors can also suppress intellectual development. As an example, early struggles to read by children with less advantageous genotypes for reading may decrease the likelihood that these children will choose reading activities, leading to slower progression of these faculties. Worse, if children’s early reading attempts are experienced very negatively, these children may develop avoidant responses to reading, which could result in active suppression of developing literacy. Importantly, when these kinds of differences originate with differences in children’s genotypes, they can increase heritability and exaggerate disparities. Identifying ECE factors that contribute to heritability of cognitive and intellectual phenotypes is important because it can point to practices that better adapt to neurogenetic diversity among children. Innovative pedagogical practices may lead to approaches that increase “enhancing” ECE effects in the subset of children disadvantaged by current practices and reduce ECE effects that suppress intellectual development and academic achievement, which may lead to more equitable educational outcomes.

### Limitations and Caveats

The proportion of variance in the cognitive measures accounted for by the genomic predictors was larger in the EurA participants than in the DivA group(Sup. Tables 5 & 7), as would be expected given the discovery samples were in individuals of European ancestry. However, the patterns were generally similar in the DivA group. This suggests similar genetic architecture for these cognitive phenotypes across ancestry groups and supports the validity of the results from the full sample. Analyses in all three groups included as covariates the top ten genetic principal components derived from the full sample. Because of broad ancestral diversity in the ABCD cohort, there is limited power for comparing the effects in different ancestry groups. As has been discussed in genetics generally(28,29), the lower predictive performance in the DivA group once again underscores the importance of collecting genetic data from ancestrally diverse populations and developing methods that can be used across ancestry groups.

One may have predicted that EAPS would have been a more powerful predictor of cognitive measures in ABCD than IPS, due to it having over 4 times the discovery sample size. However, we found generally the IPS had stronger associations (Sup. Tables 3-8), perhaps because the phenotype is a better match between training and testing. This contrasts with results of a previous study of adults, where EAPS explained 7-10% of the variance in cognition(10), while IPS explained only 2-5%(9). This discrepancy may be due to methodological differences, alternatively the young age of the cohort may be the key difference. Educational attainment, while clearly related to scores on cognitive tests, may be influenced by other genetically influenced traits (e.g. personality) that may contribute to greater persistence in formal education; thus the EAPS is likely to reflect to a greater degree these traits. Such pleiotropic of EAPS effects has been observed in adults(30). When we include both EAPS and IPS in a single model, together they explain 5.8% of the variability in the Crystallized Composite (EurA, Sup. Table 6), substantially more than IPS alone explains (4.1%), indicating that these genomic predictors capture unique sources of the relevant variance, and are likely measuring different (relevant) constructs.

These results are consistent with previous evidence for a role of genetic variation in developing cognitive functions, and they strengthen the evidence for rGE during cognitive development. However, it should be emphasized that the genomic predictors (together) account for only 4.15% of cognitive performance variance in the full sample. Furthermore, this was observed for the Crystallized Composite measure, the culturally sensitive measure hypothesized to exhibit increased genetic association as a result of rGE effects. The additive effects of potentially confounding sociocultural covariates, even controlling for IPS, accounted for 13.2% of the variability. For the Fluid Composite the genomic predictors together accounted for only 1.1% of the variance, and sociocultural covariates accounted for almost 5%. Of note, even with the narrow 2 year age range in the cohort, age alone accounts for 10% of the variability in the Crystallized Composite and 7% in the Fluid Composite. These effects may reveal clues about a highly dynamic process of cognitive and intellectual development in children.

Finally, though the results of the mediation analysis focusing on recreational reading strengthen the plausibility that such ECE mediators associate with genotypes and increase genetic effects, these results do not prove a causal explanation, and none should be inferred. In the context of an observational study such as ABCD it is always possible that confounding variables not accounted for in the analysis are responsible for the mediation effect we observed.

## ABCD Acknowledgement

Data used in the preparation of this article were obtained from the **Adolescent Brain Cognitive Development**^□^ **Study**(**ABCD Study^®^**) (https://abcdstudy.org), held in the NIMH Data Archive (NDA). This is a multisite, longitudinal study designed to recruit more than 10,000 children age 9-10 and follow them over 10 years into early adulthood. The ABCD Study is supported by the National Institutes of Health and additional federal partners under award numbers: U01DA041022, U01DA041028, U01DA041048, U01DA041089, U01DA041106, U01DA041117, U01DA041120, U01DA041134, U01DA041148, U01DA041156, U01DA041174, U24DA041123, and U24DA041147 A full list of supporters is available at https://abcdstudy.org/federal-partners/. A listing of participating sites and a complete listing of the study investigators can be found at https://abcdstudy.org/principal-investigators.html. ABCD Study consortium investigators designed and implemented the study and/or provided data but did not necessarily participate in analysis or writing of this report. This manuscript reflects the views of the authors and may not reflect the opinions or views of the NIH or ABCD Study consortium investigators. The ABCD data repository grows and changes over time. The ABCD data used in this came from [NIMH Data Archive Digital Object Identifier (10.15154/1504041)].

