## Supplementary Materials for "Gene-experience correlation during cognitive development: Evidence from the Adolescent Brain Cognitive Development (ABCD) Study^SM^"

### Supplementary Methods

#### Description of Each Cognitive Task used in ABCD

##### NIH Toolbox Cognition Battery^®^ Tasks

The *Toolbox Oral Reading Recognition Task*^®^ measured language decoding and reading. Children were asked to read aloud single letters or words presented in the center of an iPad screen. The research assistant marked pronunciations as correct or incorrect. Extensive training was given prior to administering the test battery. Item difficulty was modulated using computerized adaptive testing (CAT).

The *Toolbox Picture Vocabulary Task*^®^, a variant of the Peabody Picture Vocabulary Test (PPTV), measured language and vocabulary comprehension. Four pictures were presented on an iPad screen as a word was played through the iPad speaker. The child was instructed to point to the picture, which represented the concept, idea or object name heard. CAT was implemented to control for item difficulty and avoid floor or ceiling effects.

The *Toolbox Pattern Comparison Processing Speed Test*^®^ measured processing speed. Children were shown two images and asked to determine if they were identical or different by touching the appropriate response button on the screen. This test score is the sum of the number of items completed correctly in the time given.

The *Toolbox List Sorting Working Memory Test*^®^ measured working memory. Children heard a list of words alongside pictures of each word and were instructed to repeat the list back in order of their actual size from smallest to largest. The list started with only two items and a single category (food or animals). The number of items increased with each correct answer to a maximum of seven. The child then progressed to the next stage in which the two different categories were interleaved. At this stage children were required to report the items back in size order from the first category followed by the second category. Children were always given two opportunities to repeat the list correctly before the experimenter scored the trial as incorrect.

The *Toolbox Picture Sequence Memory Test®* measured episodic memory. On each trial, children were shown a series of fifteen pictures in a particular sequence. The pictures illustrated activities or events within a particular setting (e.g., going to the park), and as each appeared on the screen a pre-recorded narrative briefly described the content of the picture. Participants were instructed to arrange the pictures in the original sequence in which they were shown. The Rey-Auditory Verbal Learning Task was also included in the ABCD neurocognition battery as a more comprehensive measure of episodic memory.

The *Toolbox Flanker Task*^®^ measured executive function, attentional and inhibitory control. This adaptation of the Eriksen Flanker task(1) captures how readily a participant is influenced by the congruency of stimuli surrounding a target. On each trial a target arrow was presented in the center of the iPad screen facing to the left or right and was flanked by two additional arrows on both sides. The surrounding arrows were either facing in the same (congruent) or different (incongruent) direction to the central target arrow. The participant was instructed to push a response button to indicate the direction of the central target arrow. Accuracy and reaction time scores were combined to produce a total score of executive attention, such that higher scores indicate a greater ability to attend to relevant information and inhibit incorrect responses.

The *Toolbox Dimensional Change Card Sort Task*^®^ measured executive function and cognitive flexibility. On each trial, the participant was presented with two objects at the bottom of the iPad screen and a third object in the middle. The participant was asked to sort the third object by matching it to one of the bottom two objects based on either colour or shape. In the first block participants matched based on one dimension and in the second block they switched to the other dimension. In the final block, the sorting dimension alternated between trials pseudorandomly. The total score was calculated based on speed and accuracy.

##### Other Neurocognitive Tasks

*Rey-Auditory Verbal Learning Task.* This task measures auditory learning, recall and recognition. Participants listened to a list of 15 unrelated words and were asked to immediately recall these after each of five learning trials. A second unrelated list was then presented and participants were asked to recall as many words as possible from the second list and then recall words again from the initial list. Following a delay of 30 minutes (during which other non-verbal tasks from the cognitive battery are administered), longer-term retention was measured using recall and recognition. This task was administered via an iPad using the Q-interactive platform of Pearson assessments(2). In the current study, the total number of items correctly recalled across the five learning trials was summed to produce a measure of auditory verbal learning.

*Little Man Task.* This task measures visuospatial processing involving mental rotation with varying degrees of difficulty(3). A rudimentary male figure holding a briefcase in one hand was presented on an iPad screen. The figure could appear in one of four positions: right side up vs upside down and either facing the participant or with his back to the participant. The briefcase could be in either hand. Participants indicated which hand the briefcase was in using one of two buttons. Performance across the 32 trials was measured by the percentage of trials in which the child responded correctly. This was divided by the average reaction time to complete the task (in seconds) to produce a measure of efficiency of visuospatial processing. This was the dependent variable analysed for this task.

*Matrix Reasoning.* Nonverbal reasoning was measured using an automated version of the Matrix Reasoning subtest from the Weschler Intelligence Test for Children-V(4). On each trial the participant was presented with a series of visuospatial stimuli, which was incomplete. The participant was instructed to select the next stimulus in the sequence from four alternatives. There were 32 possible trials and testing ended when the participant failed three consecutive trials. The total raw score, used in the current study, was the total number of trials completed correctly.

#### Genetic Data and Computing Polygenic Scores

Saliva and blood samples were collected and sent to Rutgers University Cell and DNA Repository for DNA isolation. Genotyping was performed using the Smokescreen array(5), calling 646,247 genetic variants. Pre-variant imputation, quality control (QC) on the genotyping was performed to ensure each genetic variant had been successfully called in more than 95% of the sample, and that missingness for each individual was lower than 20%. After QC, 517,724 SNPs and 10,659 individuals remained. Based on genotype data, we derived genetic ancestry using fastStructure(6) with four ancestry groups. Genetic principal components were also calculated using PLINK.

We performed imputation using the Michigan Imputation Server(7) with the hrc.r1.1.2016 reference panel, Eagle v2.3 phasing and multiethnic imputation. PLINK(8) was used to convert dosage files to PLINK files using a best guess threshold of 0.9 for each locus. After PLINK conversion, we used post imputation variant QCs of minor allele frequency above 5%, Hardy-Weinberg threshold of 10^-6^ and no greater than 10% missing SNPs for each individual. These QC processes resulted in 1,427,972 SNPs and 10,659 individuals remaining.

We computed polygenic scores usine PRSice(9). After variant imputation we performed clumping and pruning of SNPs with a clumping window of 250 kb and r^2^ of 0.1. SNPs from the major histone compatibility complex were removed from the analysis. This resulted in 692,685 SNPs remaining. The polygenic scores for each individual were then computed as a sum of their SNPs weighted by the variant effect size in the discovery samples(10,11), with no p-value thresholding of summary statistics. Since part of the EAPS discovery sample was 23andMe participants, only the top 10,000 SNPs were included in summary statistics for this polygenic score.

#### Statistical Models for Genomic Prediction of Behavior Measures

Associations between polygenic scores and behavior measures were assessed using Generalized Linear Mixed-Effects Models (GLMMs). In addition to the IPS or EAPS, all models included the fixed effects of age, sex at birth, parental marital status, education level of parent/caregiver, household income, ethnicity (Hispanic/non-Hispanic) and the top ten genetic principal components. Data collection site and family were included as random effects. To assess variance explained by predictor(s) we computed either from t statistics and degrees of freedom (i.e. $R^{2}=t^{2}/(t^{2}+df))$ or as the computed change in *R^2^* based on a log likelihood ratio test between a full and reduced model.

#### Differential Mediation Analysis

Due to the computational burden of fitting GLMMs for bootstrapped estimates we instead fit general linear models (GLMs) with the same fixed effects as above and adding study site. To control for family we restricted this analysis to randomly selected singletons (one member from each family). By randomly selecting singletons we believe we obtain a conservative estimate of associations as we lose power when compared to GLMMs (where all family members are retained) whilst controlling family relatedness in ABCD.

|  | **Factor 1** | **Factor 2** | **Factor 3** |
| --- | --- | --- | --- |
| **Reading** | 0.82 | 0.12 | 0.12 |
| **Picture Vocabulary** | 0.75 | 0.07 | 0.19 |
| **Pattern** | 0.02 | 0.81 | 0.09 |
| **List** | 0.47 | 0.15 | 0.49 |
| **Picture** | 0.01 | 0.14 | 0.86 |
| **Flanker** | 0.21 | 0.71 | 0.07 |
| **Card Sort** | 0.21 | 0.71 | 0.23 |
| **Rey Auditory Verbal** | 0.31 | 0.13 | 0.71 |
| **Little Man Task** | 0.50 | 0.30 | 0.07 |

Sup. Figure 1 Loadings of Bayesian factors from (12), mean of posterior distributions.

| Assessment | Variables Analyzed | NDA Data Dictionary Name | Informant |
| --- | --- | --- | --- |
| NIH Toolbox® | Crystallized Composite  Fluid Composite  Reading  Picture Vocabulary  Pattern  List  Picture  Flanker  Cardsort | nihtbx_cryst_uncorrected  nihtbx_fluidcomp_uncorrected  nihtbx_reading_uncorrected  nihtbx_picvocab_uncorrected  nihtbx_pattern_uncorrected  nihtbx_pattern_uncorrected  nihtbx_list_uncorrected  nihtbx_flanker_uncorrected  nihtbx_cardsort_uncorrected | Youth |
| Little Man Task | Percentage correct | lmt_scr_perc_correct | Youth |
| Pearson Scores | Rey Auditory Verbal total correct  WISC-V Matrix Reasoning | pea_ravlt_sd_trial_[i-v]_tc (sum i-v)  pea_wiscv_trs | Youth |
| Bayesian Latent Factors | Bayesian Factor 1  Bayesian Factor 2  Bayesian Factor 3 | neurocog_pc1.bl  neurocog_pc2.bl  neurocog_pc3.bl | Youth (derived) |
| Sports and Activity Involvement Questionaries | Reading hours | sports_activity_read_hours_p | Caregiver |

Sup. Table 1 Variables used for associations, with NDA (National Institute of Mental Health Data Archive) data dictionary names.

|  | Mean (SD) | Median | Sex | Age | Sociocultural |
| --- | --- | --- | --- | --- | --- |
| Crystallized Composite | 86.87 (6.93) | 87 | 0.01 | 9.96 | 13.06 |
| Fluid Composite | 92.18 (10.43) | 93 | 0.33 | 7.32 | 4.97 |
| Reading | 91.23 (6.73) | 91 | 0.01 | 6.27 | 9.24 |
| Picture Vocabulary | 85.04 (8.02) | 84 | 0.07 | 7.94 | 10.67 |
| Pattern | 88.29 (14.47) | 88 | 0.57 | 4.85 | 1.09 |
| List | 97.43 (11.81) | 97 | 0.13 | 2.11 | 4.82 |
| Picture | 103.33 (12.01) | 103 | 0.51 | 1.20 | 2.26 |
| Flanker | 94.42 (8.83) | 96 | 0.03 | 3.26 | 2.13 |
| Cardsort | 92.97 (9.26) | 94 | 0.48 | 3.80 | 2.54 |
| Rey Auditory Verbal | 43.78 (9.96) | 44 | 1.26 | 2.32 | 3.34 |
| Matrix Reasoning | 18.13 (3.74) | 18 | 0.34 | 2.83 | 4.54 |
| Little Man Task | 0.60 (0.17) | 0.56 | 0.48 | 5.23 | 3.80 |
| Bayesian Factor 1 | 0.05 (0.76) | 0.06 | 0.30 | 10.05 | 12.73 |
| Bayesian Factor 2 | 0.02 (0.76) | 0.06 | 0.22 | 5.53 | 1.50 |
| Bayesian Factor 3 | 0.04 (0.70) | 0.04 | 0.90 | 1.64 | 2.94 |
| Recreational Reading (hours) | 6.5 (10) | 4 | 0.45 | 0.21 | 1.65 |

Sup. Table 2 Mean (SD) and median for each behavioral measure in the full sample, estimated % variance explained by sex, age, and the set of socio-cultural covariates (parental marital status, parental education, household income, genetic ancestry PCs and Hispanic/non-Hispanic), **after accounting for IPS**.

### Simulating Noise on Crystallized Composite Score

A previous study noted that crystallized composite of NIH toolbox shows greater test-retest reliability than fluid domains of intelligence(13). This is important to consider in the current study as it is possible that the observed findings, of IPS being stronger predictors of crystallized domains of intelligence, may be due differences in noise between fluid and crystallized measures. To test the robustness of the results to different degrees of noise we simulated additive noise on the crystallized composite score as:

$${Cryst}_{noise}=Cryst+N(0, \sigma)$$

In which we created samples of *Cryst_noise_* for different values of noise, $\sigma$ and repeated the main behavioral analyses with IPS.

#### IPS Predictive Performance on Cryst_nosie_ vs Fluid Composite

We simulated 1000 samples of Cryst_noise_ at each of 9 different value of $\sigma$ from 0 to 2 in increments of 0.25 resulting in 9x1000=9000 samples of Cryst_noise_. Next we performed the same z-test on performed in the main analysis, now between the standardized regression coefficients of *Cryst_noise_* and Fluid composite. We calculated the proportion of noise in signal for Cryst_noise_ as 1-r^2^, where r = cor(Cryst, Cryst_noise_). Supp. Figure 2 shows the negative log(p) value of the z-test as a function of noise, with the red dotted line indicating a significance threshold of 0.05. A previous validation study estimated the test-retest reliability of Fluid to be 0.76 and Crystallized to be 0.85(13), this means *Cryst_noise_* would need to have an r ≈ 0.76/0.85=0.89 with *Cryst* or 1-r^2^ ≈ 0.2. At this level of noise we have 1.0 power (alpha=0.05) to detect *Cryst_noise_* having a significantly greater IPS standardized regression coefficients than Fluid composite. This demonstrates that our findings are robust to the difference in noise between crystallized and fluid composite measures.

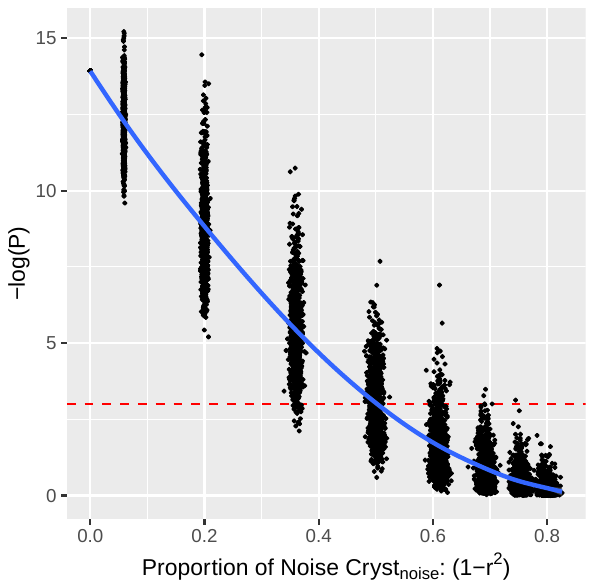

Supp. Figure 3 Significance of difference in IPS standardized regression coefficient between predicting i) Cryst_noise_ and ii) Fluid composite (y axis) as in main figure 2, as a function of proportion of noise in Cryst vs Cryst_noise_ (x-axis). Red dotted line indicates alpha=0.05.

#### Differential Mediation of Reading Between i) IPS and Cryst_nosie_ and ii) IPS and Fluid Composite

To test the robustness of the mediation results in the main analysis, we repeated the same process with 100 samples of Cryst_noise_ at each of 10 different value of $\sigma$ (0, 0.5, 0.75, 1, 1.25, 1.5, 2, 2.5, 3, 4) resulting in 1,000 different samples of Cryst_noise_. Repeating a similar mediation analysis to the main text except instead using the more classical framework of Baron and Kenny(14), due to computational feasibility, which we have found to be convergent with the mediation model(15) presented in the main text. We again looked at 5,210 singletons from the full sample to test the differential mediation of reading between IPS and Cryst_noise_ vs Fluid composite using 10,000 bootstrap samples for each Cryst_noise_, we found once again a similar robustness to the level of noise simulated. Supp. Figure 3 shows the negative log(p) for the same test performed for the difference in distributions shown in Figure 2 at the 1,000 different simulated samples of Cyrst_noise_ as a function of noise (1-r^2^). We see that when Cryst_noise_ has a value of noise making it comparable to the test retest reliability of fluid composite (1-r^2^ ≈ 0.2), all the samples remain significant. Again these simulated noise analyses demonstrate that our main results are robust to a higher degree of noise than the observed difference in test-retest reliability between Crystallized and Fluid composite measures in the toolbox.
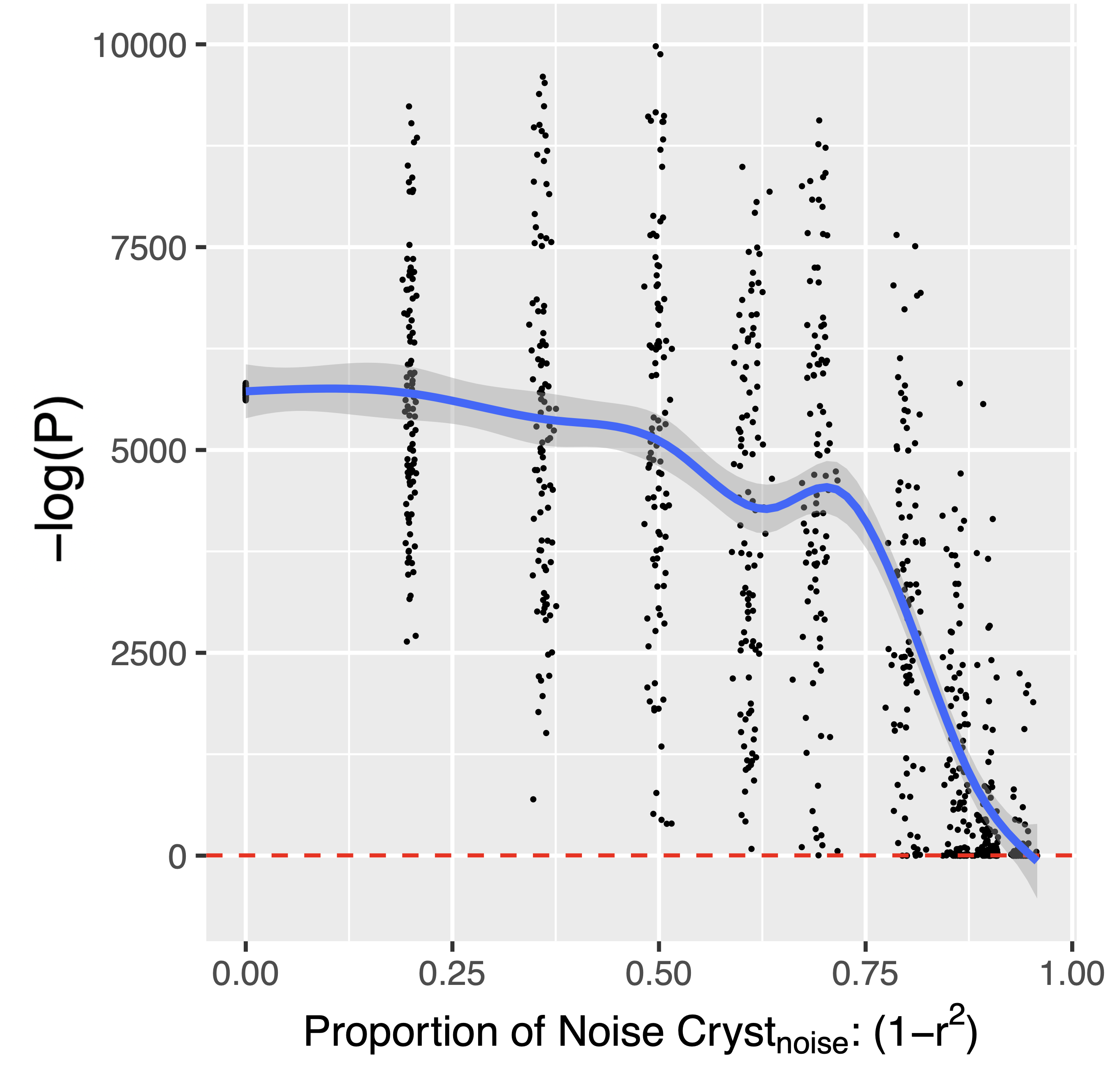

Supp. Figure 4 Welch t-test for differential mediation between i) IPS – reading - Fluid and ii) IPS – reading Cryst_noise_ (y-axis), as a function of proportion of noise in Cryst vs Cryst_noise_ (x-axis). Red dotted line indicates alpha=0.05.

### IPS, EAPS and EAPS + IPS Cognition Associations

|  | IPS | | | | EAPS | | | |
| --- | --- | --- | --- | --- | --- | --- | --- | --- |
| Behavior | coeff | Var. Explained (%) | t | pval | coeff | Var. Explained (%) | t | pval |
| Crystallized Composite | 0.50 | 2.86 | 15.82 | 1.31E-55 | 0.19 | 2.32 | 14.21 | 2.56E-45 |
| Fluid Composite | 0.28 | 0.75 | 8.03 | 1.14E-15 | 0.11 | 0.61 | 7.23 | 5.26E-13 |
| Reading | 0.49 | 2.34 | 14.27 | 1.12E-45 | 0.16 | 1.39 | 10.95 | 9.85E-28 |
| Picture Vocabulary | 0.40 | 1.79 | 12.46 | 2.59E-35 | 0.17 | 1.87 | 12.74 | 7.75E-37 |
| Pattern | 0.13 | 0.15 | 3.61 | 3.10E-04 | 0.05 | 0.15 | 3.51 | 4.44E-04 |
| List | 0.31 | 0.85 | 8.55 | 1.50E-17 | 0.11 | 0.63 | 7.33 | 2.47E-13 |
| Picture | 0.18 | 0.27 | 4.79 | 1.73E-06 | 0.09 | 0.41 | 5.88 | 4.30E-09 |
| Flanker | 0.16 | 0.23 | 4.45 | 8.70E-06 | 0.03 | 0.04 | 1.91 | 5.63E-02 |
| Cardsort | 0.14 | 0.17 | 3.77 | 1.67E-04 | 0.06 | 0.18 | 3.96 | 7.70E-05 |
| Rey Auditory Verbal | 0.32 | 0.91 | 8.81 | 1.50E-18 | 0.11 | 0.66 | 7.52 | 6.23E-14 |
| Matrix Reasoning | 0.30 | 0.84 | 8.49 | 2.34E-17 | 0.14 | 0.98 | 9.18 | 5.36E-20 |
| Little Man Task | 0.25 | 0.57 | 7.00 | 2.69E-12 | 0.07 | 0.24 | 4.55 | 5.50E-06 |
| Bayesian Factor 1 | 0.49 | 2.88 | 15.88 | 5.82E-56 | 0.17 | 1.93 | 12.92 | 7.47E-38 |
| Bayesian Factor 2 | 0.11 | 0.12 | 3.15 | 1.62E-03 | 0.03 | 0.04 | 1.80 | 7.19E-02 |
| Bayesian Factor 3 | 0.23 | 0.52 | 6.64 | 3.37E-11 | 0.11 | 0.67 | 7.55 | 4.80E-14 |

Sup. Table 3 Full sample: behavioral associations between polygenic scores i) IPS and ii) EAPS in two separate regression for each behavior using GLMMs and controlling for variables of no interest.

| Behavior | EAPS + IPS effect | | | EAPS effect beyond IPS alone | | |
| --- | --- | --- | --- | --- | --- | --- |
|  | Var. Explained (%) | $\boldsymbol{\chi}^{\boldsymbol{2}}$ | pval | Var. Explained  (%) | $\boldsymbol{\chi}^{\boldsymbol{2}}$ | pval |
| Crystallized Composite | 4.15 | 360.91 | 4.26E-79 | 1.33 | 113.72 | 1.50E-26 |
| Fluid Composite | 1.09 | 93.46 | 5.08E-21 | 0.34 | 29.09 | 6.90E-08 |
| Reading | 3.01 | 260.70 | 2.46E-57 | 0.69 | 58.99 | 1.59E-14 |
| Picture Vocabulary | 2.93 | 253.05 | 1.12E-55 | 1.16 | 99.06 | 2.44E-23 |
| Pattern | 0.24 | 20.28 | 3.95E-05 | 0.08 | 7.25 | 7.09E-03 |
| List | 1.19 | 101.62 | 8.56E-23 | 0.34 | 28.68 | 8.54E-08 |
| Picture | 0.54 | 46.18 | 9.37E-11 | 0.27 | 23.25 | 1.42E-06 |
| Flanker | 0.24 | 20.50 | 3.54E-05 | 0.01 | 0.66 | 4.17E-01 |
| Cardsort | 0.28 | 23.91 | 6.44E-06 | 0.11 | 9.67 | 1.87E-03 |
| Rey Auditory Verbal | 1.25 | 107.51 | 4.51E-24 | 0.35 | 30.02 | 4.27E-08 |
| Matrix Reasoning | 1.46 | 124.94 | 7.40E-28 | 0.62 | 52.99 | 3.36E-13 |
| Little Man Task | 0.67 | 57.33 | 3.56E-13 | 0.10 | 8.26 | 4.05E-03 |
| Bayesian Factor 1 | 3.88 | 337.50 | 5.16E-74 | 1.04 | 88.74 | 4.50E-21 |
| Bayesian Factor 2 | 0.13 | 11.07 | 3.94E-03 | 0.01 | 1.11 | 2.91E-01 |
| Bayesian Factor 3 | 0.95 | 81.48 | 2.03E-18 | 0.44 | 37.44 | 9.44E-10 |

Sup. Table 4 Full sample: Combined (EAPS + IPS ) regression on behavior. Left 3 columns show effect of EAPS + IPS, right 3 columns show EAPS effect after controlling for IPS.

|  | IPS | | | | EAPS | | | |
| --- | --- | --- | --- | --- | --- | --- | --- | --- |
| Behavior | coeff | Var. Explained (%) | t | pval | coeff | Var. Explained (%) | t | pval |
| Crystallized Composite | 0.21 | 4.13 | 14.48 | 1.44E-46 | 0.18 | 3.34 | 12.95 | 9.36E-38 |
| Fluid Composite | 0.11 | 1.15 | 7.53 | 6.10E-14 | 0.09 | 0.89 | 6.60 | 4.66E-11 |
| Reading | 0.20 | 3.49 | 13.25 | 2.05E-39 | 0.14 | 2.03 | 10.04 | 1.64E-23 |
| Picture Vocabulary | 0.16 | 2.56 | 11.30 | 2.90E-29 | 0.16 | 2.71 | 11.64 | 6.71E-31 |
| Pattern | 0.05 | 0.24 | 3.41 | 6.65E-04 | 0.04 | 0.18 | 2.94 | 3.27E-03 |
| List | 0.12 | 1.21 | 7.72 | 1.41E-14 | 0.10 | 0.95 | 6.83 | 9.50E-12 |
| Picture | 0.07 | 0.38 | 4.29 | 1.84E-05 | 0.07 | 0.50 | 4.96 | 7.42E-07 |
| Flanker | 0.08 | 0.50 | 4.95 | 7.72E-07 | 0.04 | 0.12 | 2.37 | 1.80E-02 |
| Cardsort | 0.06 | 0.27 | 3.64 | 2.74E-04 | 0.05 | 0.25 | 3.49 | 4.83E-04 |
| Rey Auditory Verbal | 0.11 | 1.13 | 7.44 | 1.17E-13 | 0.10 | 1.03 | 7.11 | 1.32E-12 |
| Matrix Reasoning | 0.11 | 1.10 | 7.35 | 2.24E-13 | 0.11 | 1.08 | 7.27 | 4.10E-13 |
| Little Man Task | 0.09 | 0.74 | 6.03 | 1.75E-09 | 0.05 | 0.22 | 3.31 | 9.45E-04 |
| Bayesian Factor 1 | 0.20 | 4.12 | 14.46 | 1.88E-46 | 0.16 | 2.75 | 11.73 | 2.43E-31 |
| Bayesian Factor 2 | 0.05 | 0.22 | 3.27 | 1.09E-03 | 0.02 | 0.06 | 1.65 | 9.89E-02 |
| Bayesian Factor 3 | 0.08 | 0.64 | 5.61 | 2.10E-08 | 0.10 | 0.90 | 6.64 | 3.37E-11 |

Sup. Table 5 European ancestry sample: behavioral associations between polygenic scores i) IPS and ii) EAPS in two separate regression for each behavior using GLMMs and controlling for variables of no interest.

| Behavior | EAPS + IPS effect | | | EAPS effect beyond IPS alone | | |
| --- | --- | --- | --- | --- | --- | --- |
|  | Var. Explained (%) | $\boldsymbol{\chi}^{\boldsymbol{2}}$ | pval | Var. Explained  (%) | $\boldsymbol{\chi}^{\boldsymbol{2}}$ | pval |
| Crystallized Composite | 5.80 | 291.74 | 4.46E-64 | 1.74 | 85.82 | 1.97E-20 |
| Fluid Composite | 1.58 | 78.02 | 1.15E-17 | 0.44 | 21.38 | 3.76E-06 |
| Reading | 4.33 | 216.21 | 1.12E-47 | 0.88 | 43.24 | 4.85E-11 |
| Picture Vocabulary | 4.08 | 203.45 | 6.62E-45 | 1.56 | 76.94 | 1.76E-18 |
| Pattern | 0.32 | 15.78 | 3.74E-04 | 0.08 | 4.17 | 4.11E-02 |
| List | 1.68 | 82.56 | 1.18E-18 | 0.47 | 23.08 | 1.55E-06 |
| Picture | 0.68 | 33.48 | 5.37E-08 | 0.30 | 14.96 | 1.10E-04 |
| Flanker | 0.52 | 25.47 | 2.94E-06 | 0.02 | 0.95 | 3.29E-01 |
| Cardsort | 0.40 | 19.76 | 5.13E-05 | 0.13 | 6.46 | 1.10E-02 |
| Rey Auditory Verbal | 1.66 | 82.04 | 1.53E-18 | 0.54 | 26.67 | 2.41E-07 |
| Matrix Reasoning | 1.68 | 82.92 | 9.89E-19 | 0.59 | 28.93 | 7.51E-08 |
| Little Man Task | 0.80 | 39.07 | 3.28E-09 | 0.05 | 2.64 | 1.04E-01 |
| Bayesian Factor 1 | 5.39 | 270.80 | 1.57E-59 | 1.33 | 65.36 | 6.25E-16 |
| Bayesian Factor 2 | 0.23 | 11.28 | 3.55E-03 | 0.01 | 0.56 | 4.53E-01 |
| Bayesian Factor 3 | 1.21 | 59.41 | 1.26E-13 | 0.57 | 27.83 | 1.32E-07 |

Sup. Table 6 European ancestry sample: Combined (EAPS + IPS ) regression on behavior. Left 3 columns show effect of EAPS + IPS, right 3 columns show EAPS effect after controlling for IPS.

|  | IPS | | | | EAPS | | | |
| --- | --- | --- | --- | --- | --- | --- | --- | --- |
| Behavior | coeff | Var. Explained (%) | t | pval | coeff | Var. Explained (%) | t | pval |
| Crystallized Composite | 0.40 | 1.47 | 7.34 | 2.68E-13 | 0.15 | 1.21 | 6.66 | 3.24E-11 |
| Fluid Composite | 0.20 | 0.32 | 3.41 | 6.52E-04 | 0.08 | 0.32 | 3.38 | 7.28E-04 |
| Reading | 0.39 | 1.21 | 6.64 | 3.61E-11 | 0.12 | 0.66 | 4.90 | 1.01E-06 |
| Picture Vocabulary | 0.32 | 0.89 | 5.70 | 1.30E-08 | 0.14 | 1.02 | 6.11 | 1.13E-09 |
| Pattern | 0.10 | 0.07 | 1.58 | 1.14E-01 | 0.05 | 0.10 | 1.88 | 6.08E-02 |
| List | 0.26 | 0.47 | 4.13 | 3.64E-05 | 0.09 | 0.30 | 3.30 | 9.87E-04 |
| Picture | 0.12 | 0.09 | 1.83 | 6.76E-02 | 0.08 | 0.25 | 2.98 | 2.92E-03 |
| Flanker | 0.09 | 0.05 | 1.39 | 1.64E-01 | 0.01 | 0.01 | 0.44 | 6.58E-01 |
| Cardsort | 0.10 | 0.07 | 1.59 | 1.13E-01 | 0.06 | 0.13 | 2.19 | 2.83E-02 |
| Rey Auditory Verbal | 0.28 | 0.56 | 4.53 | 6.13E-06 | 0.08 | 0.23 | 2.90 | 3.70E-03 |
| Matrix Reasoning | 0.26 | 0.50 | 4.28 | 1.93E-05 | 0.14 | 0.87 | 5.62 | 2.07E-08 |
| Little Man Task | 0.21 | 0.31 | 3.37 | 7.64E-04 | 0.08 | 0.24 | 2.97 | 2.96E-03 |
| Bayesian Factor 1 | 0.40 | 1.57 | 7.59 | 4.10E-14 | 0.14 | 1.03 | 6.12 | 1.04E-09 |
| Bayesian Factor 2 | 0.06 | 0.03 | 0.96 | 3.39E-01 | 0.02 | 0.02 | 0.74 | 4.57E-01 |
| Bayesian Factor 3 | 0.19 | 0.27 | 3.14 | 1.73E-03 | 0.09 | 0.32 | 3.39 | 7.03E-04 |

Sup. Table 7 Diverse ancestry sample: behavioral associations between polygenic scores i) IPS and ii) EAPS in two separate regression for each behavior using GLMMs and controlling for variables of no interest.

| Behavior | EAPS + IPS effect | | | EAPS effect beyond IPS alone | | |
| --- | --- | --- | --- | --- | --- | --- |
|  | Var. Explained (%) | $\boldsymbol{\chi}^{\boldsymbol{2}}$ | pval | Var. Explained  (%) | $\boldsymbol{\chi}^{\boldsymbol{2}}$ | pval |
| Crystallized Composite | 2.27 | 83.46 | 7.54E-19 | 0.81 | 29.70 | 5.04E-08 |
| Fluid Composite | 0.54 | 19.64 | 5.44E-05 | 0.22 | 7.94 | 4.84E-03 |
| Reading | 1.60 | 58.42 | 2.06E-13 | 0.39 | 14.33 | 1.53E-04 |
| Picture Vocabulary | 1.62 | 59.33 | 1.31E-13 | 0.73 | 26.79 | 2.27E-07 |
| Pattern | 0.14 | 5.16 | 7.56E-02 | 0.07 | 2.65 | 1.03E-01 |
| List | 0.65 | 23.75 | 6.95E-06 | 0.19 | 6.64 | 9.97E-03 |
| Picture | 0.29 | 10.56 | 5.10E-03 | 0.20 | 7.24 | 7.12E-03 |
| Flanker | 0.05 | 1.99 | 3.69E-01 | 0.00 | 0.04 | 8.43E-01 |
| Cardsort | 0.17 | 6.35 | 4.17E-02 | 0.10 | 3.80 | 5.12E-02 |
| Rey Auditory Verbal | 0.69 | 25.04 | 3.65E-06 | 0.12 | 4.51 | 3.36E-02 |
| Matrix Reasoning | 1.17 | 42.75 | 5.22E-10 | 0.67 | 24.39 | 7.86E-07 |
| Little Man Task | 0.47 | 17.19 | 1.85E-04 | 0.16 | 5.79 | 1.61E-02 |
| Bayesian Factor 1 | 2.21 | 81.27 | 2.25E-18 | 0.66 | 23.84 | 1.05E-06 |
| Bayesian Factor 2 | 0.03 | 1.27 | 5.30E-01 | 0.01 | 0.34 | 5.57E-01 |
| Bayesian Factor 3 | 0.50 | 18.16 | 1.14E-04 | 0.23 | 8.35 | 3.86E-03 |

Sup. Table 8 Diverse ancestry sample: Combined (EAPS + IPS ) regression on behavior. Left 3 columns show effect of EAPS + IPS, right 3 columns show EAPS effect after controlling for IPS.

1.	Eriksen BA, Eriksen CW. Effects of noise letters upon the identification of a target letter in a nonsearch task. Percept Psychophys. 1974 Jan;16(1):143–9.

2.	Daniel MH, Wahlstrom D, Zhang O. Equivalence of Q-interactive^TM^ and Paper Administrations of Cognitive Tasks: WISC ®-V Q-interactive Technical Report 8. 2014.

3.	Acker W. A computerized approach to psychological screening - The Bexley-Maudsley Automated Psychological Screening and The Bexley-Maudsley Category Sorting Test. Int J Human-computer Stud \/ Int J Man-machine Stud. 1982;17:361–9.

4.	Weschler D. Weschler Intelligence Scale for Children, 5th ed. V. Bloomington, MN: Pearson; 2014.

11.	Zacher M, Nguyen-viet TA, Bowers P, Sidorenko J, Linnér RK. Gene discovery and polygenic prediction from a genome-wide association study of educational attainment in 1.1 million individuals. 2018;50(August).

12.	Thompson WK, Barch DM, Bjork JM, Gonzalez R, Nagel BJ, Nixon SJ, et al. The structure of cognition in 9 and 10 year-old children and associations with problem behaviors: Findings from the ABCD study’s baseline neurocognitive battery. Dev Cogn Neurosci [Internet]. 2018;(December):100606. Available from: https://doi.org/10.1016/j.dcn.2018.12.004

13.	Akshoomoff N, Beaumont JL, Bauer PJ, Dikmen S. NIH Toolbox Cognitive Function Battery (CFB): Composite Scores of Crystallized, Fluid, and Overall Cognition. Monogr Soc Res Child Dev. 2013;78(4):119–32.

14.	Baron R, Kenny D. The Moderator-Mediator Variable Distinction in Social Psychological Research: Conceptual, Strategic, and Statistical Considerations. J Peronality Soc Psychol. 1986;51(6):1173–82.

15.	Tingley D, Yamamoto T, Hirose K, Keele L, Imai K. Mediation: R package for causal mediation analysis. J Stat Softw. 2014;59(5):1–38.
